# Acute Social Stress Engages Synergistic Activity of Stress Mediators in the VTA to Promote Pavlovian Reward Learning

**DOI:** 10.1101/191148

**Authors:** Jorge Tovar-Diaz, Matthew B. Pomrenze, Bahram Pahlavan, Russell Kan, Michael R. Drew, Hitoshi Morikawa

## Abstract

Stressful events rapidly trigger activity-dependent synaptic plasticity in certain brain areas, driving the formation of aversive memories. However, it remains unclear how stressful experience affects plasticity mechanisms to regulate learning of appetitive events, such as intake of addictive drugs or palatable foods. Using rats, we show that two acute stress mediators, corticotropin-releasing factor (CRF) and norepinephrine (NE), enhance plasticity of NMDA receptor-mediated glutamatergic transmission in the ventral tegmental area (VTA) through their differential effects on inositol 1,4,5-triphosphate (IP)-dependent Ca^2+^ signaling. In line with this, acute social defeat stress engages convergent CRF and NE signaling in the VTA to enhance learning of cocaine-paired cues. Furthermore, defeat stress enables learning of a food-paired cue with no delay between the cue onset and food delivery. We propose that acute stress mediators synergistically regulate IP_3_-Ca^2+^ signaling in the VTA to promote appetitive Pavlovian conditioning, likely enabling learning of cues with no predictive value.

## INTRODUCTION

Stressor intensity, controllability, and duration are major determinants for regulation of future stress coping behavior and diverse cognitive functions (Koolhaas et al., 2011). In general, acute mild-to-moderate stress energizes adaptive cognitive processes and behaviors in the short run while severe/uncontrollable/chronic stress leads to maladaptive changes in brain function, including hippocampus-dependent learning and memory and other higher order cognitive processes (Chattarji et al., 2015; Kim et al., 2015; McEwen, 2007). As these cognitive functions are primarily declarative and studied outside the context of emotional valence, less is known about the impact of stress on reward-driven learning and behavior [see (Rodrigues et al., 2009) for stress effect on fear learning]. In this regard, stress is a well-known risk factor for the development of addiction, which can be viewed as a maladaptive form of reward learning (Sinha, 2008). While many studies have linked stress to addiction through long-term influence of glucocorticoids in the brain, stress can also exert rapid effects through the release of corticotropin-releasing factor (CRF) and norepinephrine (NE) (Joels et al., 2011; Maras and Baram, 2012). Immediate impact of stress has been studied extensively in intensification and/or reinstatement of drug seeking (Mantsch et al., 2016; Polter and Kauer, 2014); however it is not clear how stressful experience acutely regulates the acquisition of addictive behavior.

Dopamine (DA) neurons in the ventral tegmental area (VTA) play a key role in reward learning (Schultz, 2015). These neurons display transient burst firing in response to primary rewards (e.g., palatable food), while addictive drugs induce repetitive DA neuron bursting via pharmacological actions (Covey et al., 2014; Keiflin and Janak,2015). During cue-reward conditioning, DA neurons “learn” to respond to reward-predicting cues, thereby encoding the positive emotional/motivational valence of those cues (Cohen et al., 2012; Schultz, 1998; Stauffer et al., 2016). Glutamatergic inputs onto DA neurons drive burst firing via activation of NMDA receptors (Overton and Clark, 1997; Paladini and Roeper, 2014); thus strengthening of cue-driven NMDA input may contribute to conditioned bursting. We have shown previously that repeated pairing of cue-like glutamatergic input stimulation with reward-like bursting leads to long-term potentiation (LTP) of NMDA transmission (LTP-NMDA) in DA neurons (Harnett et al., 2009). LTP induction requires amplification of burst-evoked Ca^2+^ signals by preceding activation of metabotropic glutamate receptors (mGluRs) coupled to the generation of inositol 1,4,5-triphosphate (IP_3_). Here, IP_3_ receptors (IP_3_Rs) detect the coincidence of IP_3_ generated by glutamatergic input activity and burst-driven Ca^2+^ entry. Mechanistically, IP_3_ enhances Ca^2+^ activation of IP_3_Rs, thereby promoting Ca^2+^-induced Ca^2+^ release from intracellular stores (Taylor and Laude, 2002). In this study, we demonstrate how CRF and NE actions in the VTA regulate plasticity of NMDA transmission and the impact of acute stress on Pavlovian cue-reward learning.

## RESULTS

### Acute social stress enhances cocaine-associated cue learning

We first investigated how acute social defeat stress affects the learning of cocaine-associated cues using a conditioned place preference (CPP) paradigm. Rats underwent 30-min social defeat (∼5 min of direct contact/defeat followed by ∼25 min of protected threat), a form of uncontrollable psychosocial stress that elicits strong physiological responses (Koolhaas et al., 2011). After a 10-min interval, these stressed rats and handled controls were conditioned with a relatively low dose of cocaine (5 mg/kg, i.p.; Figure 1A). This acute defeat stress–cocaine conditioning sequence was limited to a single session to eliminate the confounding effect reflecting persistent influence of stress on CPP acquisition and/or expression (Burke et al., 2011; Chuang et al., 2011; Kreibich et al., 2009; Smith et al., 2012; Stelly et al., 2016). We found that stressed rats developed larger preference for the cocaine-paired chamber compared to control rats (Figure 1B,C,F). Both stressed and control rats developed comparable robust CPP with an increase in cocaine dose (10 mg/kg) during conditioning (Figure 1D–F). Defeat stress failed to affect CPP when cocaine conditioning (5 mg/kg) was performed after a prolonged interval (1.5 hr; Figure 1G–J). These results show that social defeat stress acutely increases the sensitivity to cocaine conditioning.

**Figure 1.**
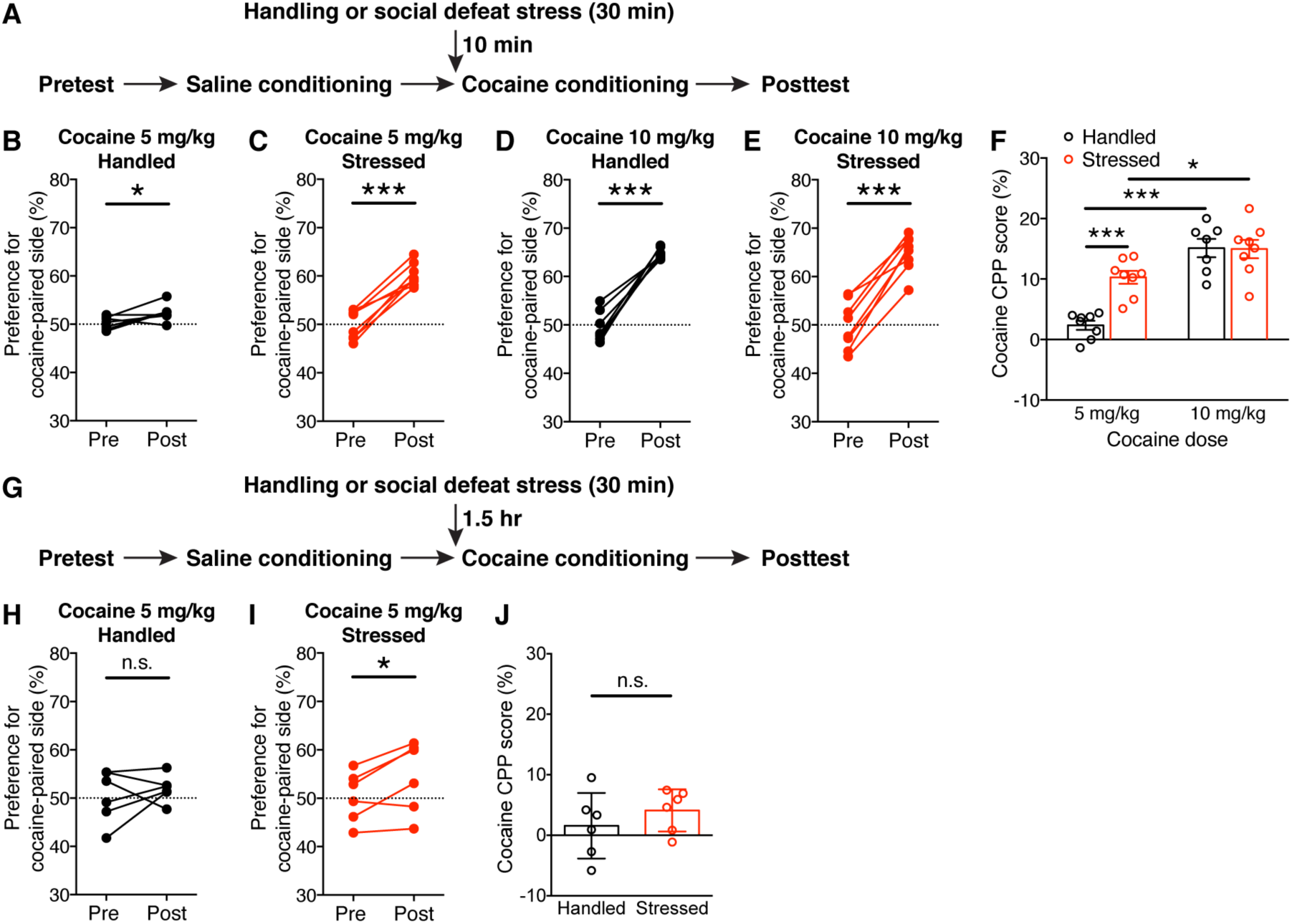
Acute exposure to social defeat stress enhances cocaine place conditioning. (A) Experimental timeline for testing the effect of acute social defeat stress on acquisition of cocaine CPP. (B–E) Changes in the preference for the cocaine-paired side in handled control rats and stressed rats conditioned with 5 mg/kg or 10 mg/kg cocaine (B: t_7_ = 3.14, p < 0.05; C: t_7_ = 9.61, p < 0.0001; D: t_6_ = 9.97, p < 0.0001; E: t_7_ = 9.82, p < 0.0001; two-tailed paired t-test). (F) Summary graph demonstrating defeat stress-induced enhancement of sensitivity to cocaine conditioning (stress: F_1,27_ = 9.81, p < 0.01; cocaine dose: F_1,27_ = 49.3, p < 0.0001; stress × cocaine dose: F_1,27_ = 10.62, p < 0.01; two-way ANOVA). *p < 0.05, ***p < 0.001 (Bonferroni post hoc test). (G) Experimental timeline for testing the effect of social defeat stress on cocaine CPP after a 1.5-hr interval. (H and I) Changes in the preference for the cocaine-paired side in rats that underwent handling (H) or social defeat (I) 1.5 hr before cocaine conditioning (5 mg/kg) (H: t_5_ = 0.71, p = 0.51; I: t_5_ = 2.8, p < 0.05; two-tailed paired t-test). (J) Graph illustrating the ineffectiveness of defeat stress on cocaine CPP with a prolonged interval (t_10_ = 0.96, p = 0.36; two-tailed unpaired t-test).

### CRF and NE differentially and synergistically promote NMDA plasticity in the VTA

Potentiation of NMDA excitation of DA neurons in the VTA may contribute to the learning of cues associated with rewards, including addictive drugs (Stelly et al., 2016; Wang et al., 2011; Whitaker et al., 2013; Zweifel et al., 2008; Zweifel et al., 2009). CRF and NE are the two major mediators of short-term stress effects in the brain (Joels et al., 2011; Maras and Baram, 2012). To gain insight into the mechanisms underlying acute defeat stress-induced enhancement of cocaine conditioning, we examined the effect of CRF and NE on NMDA plasticity using ex vivo VTA slices.

Induction of LTP-NMDA requires mGluR/IP_3_-dependent facilitation of action potential (AP)-evoked Ca^2+^ signals (Harnett et al., 2009). CRF enhances IP_3_-Ca^2+^ signaling by activation of CRF receptor 2 (CRFR2) in DA neurons (Bernier et al., 2011; Riegel and Williams, 2008; Whitaker et al., 2013), likely via protein kinase A (PKA)-mediated phosphorylation causing increased IP_3_R sensitivity (Wagner et al., 2008). To first confirm this CRF effect, we assessed AP-evoked Ca^2+^ signals using the size of Ca^2+^-sensitive K^+^ currents (I_K(Ca)_) and a subthreshold concentration of IP_3_ (10 µM·mW) was photolytically applied into the cytosol for 100 ms immediately before evoking unclamped APs (see Methods and Materials). Bath application of CRF (100 nM) significantly increased the magnitude of IP_3_-induced facilitation of I_K(Ca)_ (Figure 2A,B).

**Figure 2.**
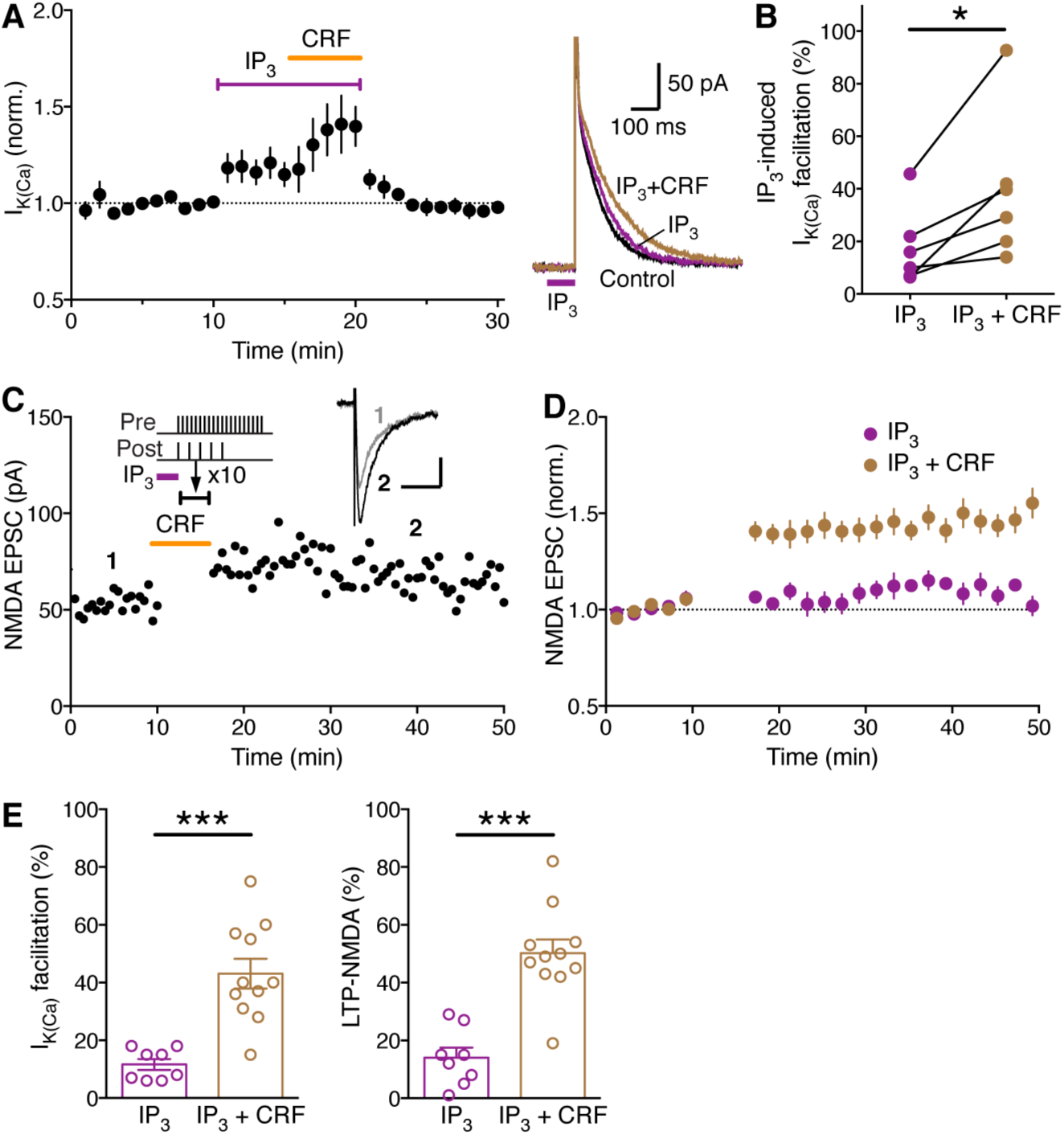
CRF enhances induction of LTP-NMDA driven by IP_3_-induced Ca^2+^ signal facilitation in VTA dopamine neurons. (A) Summary time graph (left) and example traces (right) showing that bath application of CRF (100 nM) augments IP_3_-induced facilitation of AP-evoked I_K(Ca)_. Subthreshold level of IP_3_ (determined as shown in Figure 2–figure supplement 1) was photolytically applied into the cytosol for 100 ms (purple bar in example traces) immediately before evoking unclamped APs (6 cells from 4 rats). (B) Graph plotting the magnitude of IP_3_-induced I_KCa_ facilitation before and after CRF application (t_5_ = 3.29, p < 0.05, two-tailed paired t-test). (C) Representative experiment to induce LTP in the presence of CRF. CRF (100 nM) was perfused for ∼6 min after 10-min baseline EPSC recording, while the LTP induction (A) protocol, which consisted of IP_3_-synaptic stimulation-burst combination (illustrated at the top), was delivered at the time indicated (10 times every 20 s; during a 3-min period starting ∼3 min after the onset of CRF perfusion to allow for CRF effect to take place; see panel A). Example traces of NMDA EPSCs at times indicated are shown in inset (scale bars: 50 ms/20 pA). (D) Summary time graph of LTP experiments in which LTP was induced using an IP_3_-synaptic stimulation-burst combination protocol in control solution (8 cells from 8 rats) and in CRF (11 cells from 9 rats). (E) Summary bar graphs depicting the magnitude of I_k(Ca)_ facilitation (left) and LTP (right) for the experiments shown in (D). IP_3_-induced facilitation of single AP-evoked I_K(Ca)_ was assessed by comparing the size of I_K(Ca)_ with and without preceding IP_3_ application, which was done immediately before or after delivering the LTP induction protocol (I_k(Ca)_ facilitation: t_17_ = 5.01, p < 0.0001; LTP: t_17_ = 5.70, p < 0.0001; two-tailed unpaired t-test).

Next, the effect of CRF on LTP-NMDA was tested using an induction protocol consisting of subthreshold IP_3_ application (100 ms) prior to simultaneous pairing of a burst (5 APs at 20 Hz) with a brief train of synaptic stimulation (20 stimuli at 50 Hz), the latter being necessary to induce LTP at specific inputs likely via activating NMDA receptors at those inputs at the time of burst (Harnett et al., 2009; Stelly et al., 2016; Whitaker et al., 2013). While this induction protocol using a low concentration of IP_3_ (10µM·mW) produced relatively small LTP in control solution, robust LTP was induced in the presence of CRF (100 nM; Figure 2C–E).

We further examined the effect of CRF on I_K(Ca)_ and LTP induction without IP_3_ application. CRF (100–300 nM) had a small effect on I_K(Ca)_ (Figure 3A,B), likely reflecting facilitation of small IP_3_R-mediated Ca^2+^-induced Ca^2+^ release triggered by APs themselves in DA neurons (Cui et al., 2004). Consistent with this observation, CRF failed to enable measurable LTP when simultaneous synaptic stimulation-burst pairing without prior IP_3_ application was used to induce LTP (Figure 3C–E).

**Figure 3.**
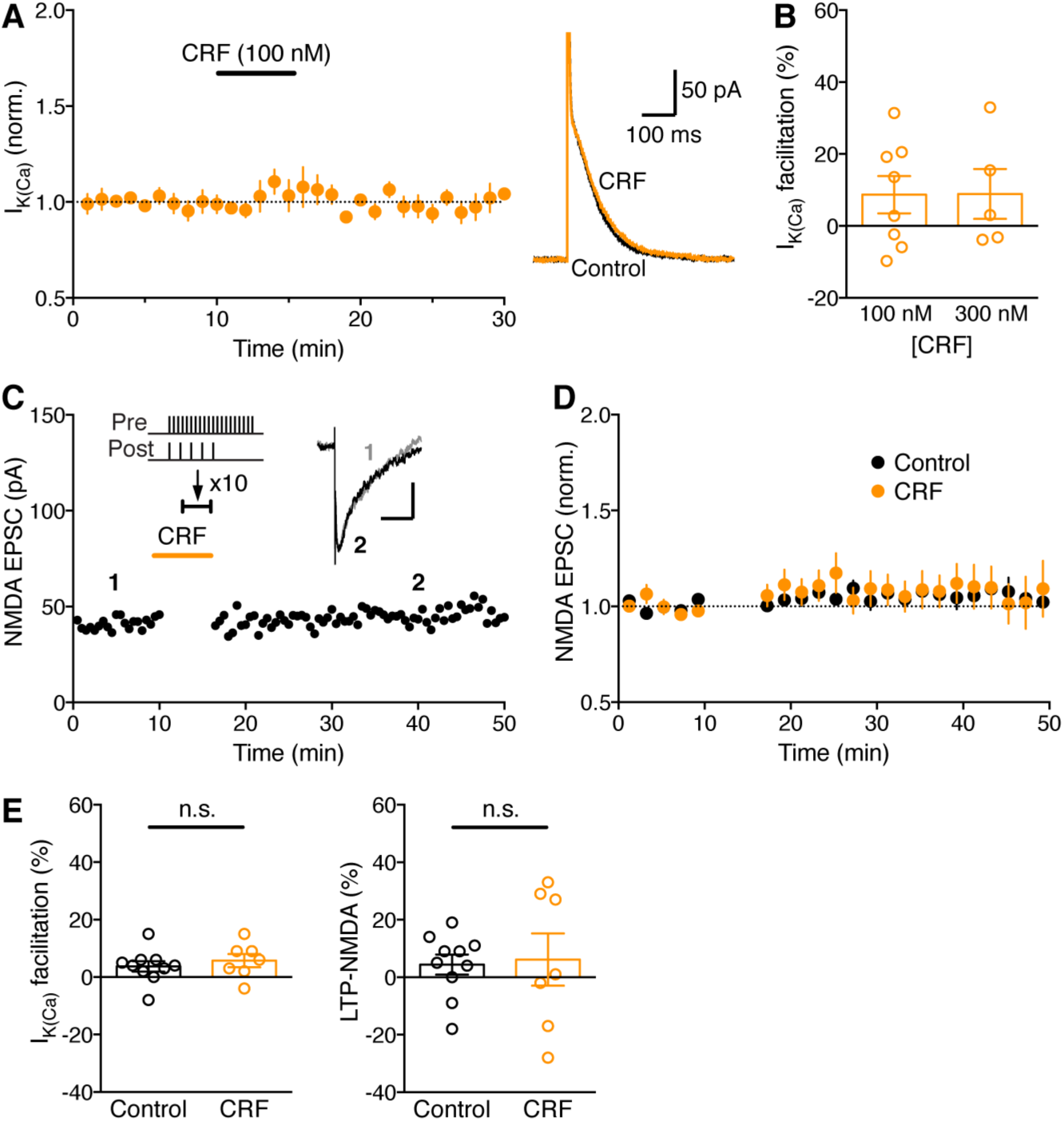
CRF causes no LTP without IP_3_-induced Ca^2+^ signal facilitation. (A) Summary time graph (left) and example traces (right) illustrating that CRF (100 nM) has small effect on I_K(Ca)_ without preceding IP_3_ application (8 cells from 7 rats). (B) Summary bar graph showing the magnitude of I_K(Ca)_ facilitation produced by two concentrations of CRF (300 nM: 5 cells from 4 rats). (C) Representative experiment to induce LTP in the presence of CRF using an induction protocol consisting of synaptic stimulation-burst pairing with no preceding IP_3_ application. Example EPSC traces at the times indicated are shown in inset (scale bars: 50 ms/20 pA). (D) Summary time graph of LTP experiments in which LTP was induced using a synaptic stimulation-burst pairing protocol in control solution (10 cells from 10 rats) and in CRF (7 cells from 7 rats). (E) Summary bar graphs depicting the magnitude of I_k(Ca)_ facilitation (left) and LTP (right) for the experiments shown in (D). I_K(Ca)_ facilitation was assessed by comparing the size of single AP-evoked I_K(Ca)_ measured immediately after 10-min baseline EPSC recording with that measured immediately before or after LTP induction (I_k(Ca)_ facilitation: t_15_ = 0.70, p = 0.50; LTP: t_15_ = 0.20, p = 0.84; two-tailed unpaired t-test).

DA neurons express α1 adrenergic receptors (α1ARs) that are coupled to phospholipase C-mediated IP_3_ synthesis (Cui et al., 2004; Paladini et al., 2001). Accordingly, bath application of the α1AR agonist phenylephrine (0.5–1 µM) increased I_K(Ca)_ in a concentration-dependent manner in the absence of exogenous IP_3_ application (Figure 4A,B). Phenylephrine treatment enabled robust LTP induction with simultaneous synaptic stimulation-burst pairing (Figure 4C–E; see Figure 4–figure supplement 1 for NE effect), in contrast to the ineffectiveness of CRF described above.

**Figure 4.**
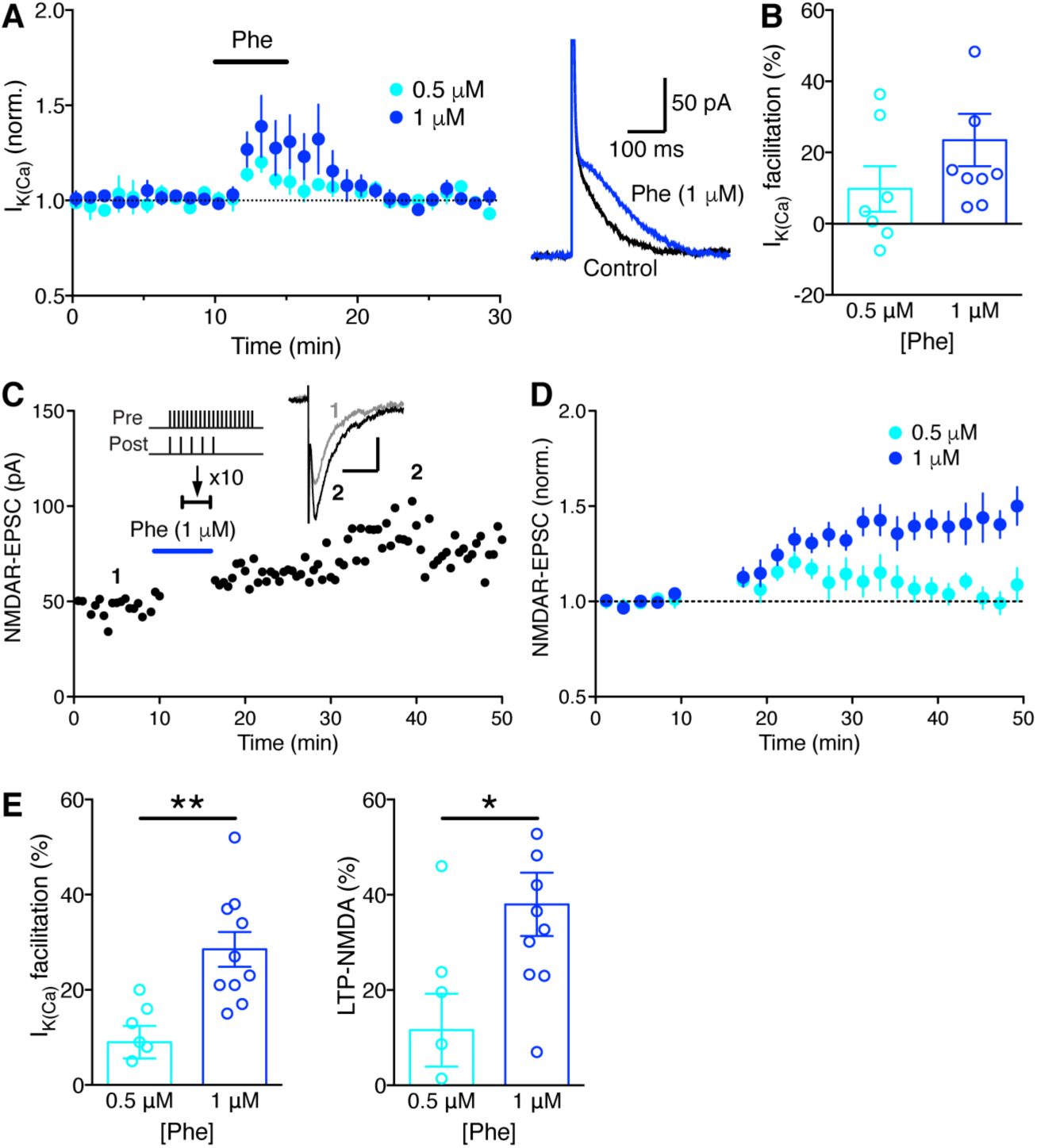
α1AR agonist phenylephrine enables LTP without IP_3_-induced Ca^2+^ signal facilitation. (A) Summary time graph (left) and example traces (right) depicting the facilitatory effect of two concentrations of phenylephrine on I_K(Ca)_ (0.5 µM: 7 cells from 3 rats; 1 µM: 9 cells from 6 rats). (B) Summary bar graph showing the magnitude of phenylephrine-induced I_K(Ca)_ facilitation. (C) Representative experiment to induce LTP-NMDA in the presence of phenylephrine (1 µM) using an induction protocol consisting of synaptic stimulation-burst pairing with (A) no preceding IP_3_ application. Example EPSC traces at the times indicated are shown in inset (scale bars: 50 ms/20 pA). (D) Summary time graph of LTP experiments in which LTP was induced using a synaptic stimulation-burst pairing protocol in the presence of 0.5 µM or 1 µM phenylephrine (0.5 µM: 7 cells from 7 rats; 1 µM: 10 cells from 8 rats). (E) Summary bar graphs depicting the magnitude of I_k(Ca)_ facilitation (left) and LTP (right) for the experiments shown in (D). I_K(Ca)_ facilitation was assessed by comparing the size of single AP-evoked I_K(Ca)_ measured immediately after 10-min baseline EPSC recording with that measured immediately before or after LTP induction (I_k(Ca)_ facilitation: t_15_ = 3.72, p < 0.01; LTP: t_15_ = 2.59, p < 0.05; two-tailed unpaired t-test).

We next asked if CRF, via CRFR2-mediated IP_3_R sensitization, could enhance the effect of phenylephrine. CRF (100 nM), which had minimal effect on I_K(Ca)_ by itself (Figure 3A,B), significantly augmented the small I_K(Ca)_ facilitation produced by a low concentration (0.5 µM) of phenylephrine (Figure 5A,B), while there was no significant CRF effect on I_K(Ca)_ facilitation caused by 1 µM phenylephrine (Figure 5–figure supplement 1). As a consequence, combined application of CRF and 0.5 µM phenylephrine enabled LTP with simultaneous synaptic stimulation-burst pairing protocol, comparable to LTP induced in the presence of 1 µM phenylephrine (Figure 5C,D).

**Figure 5.**
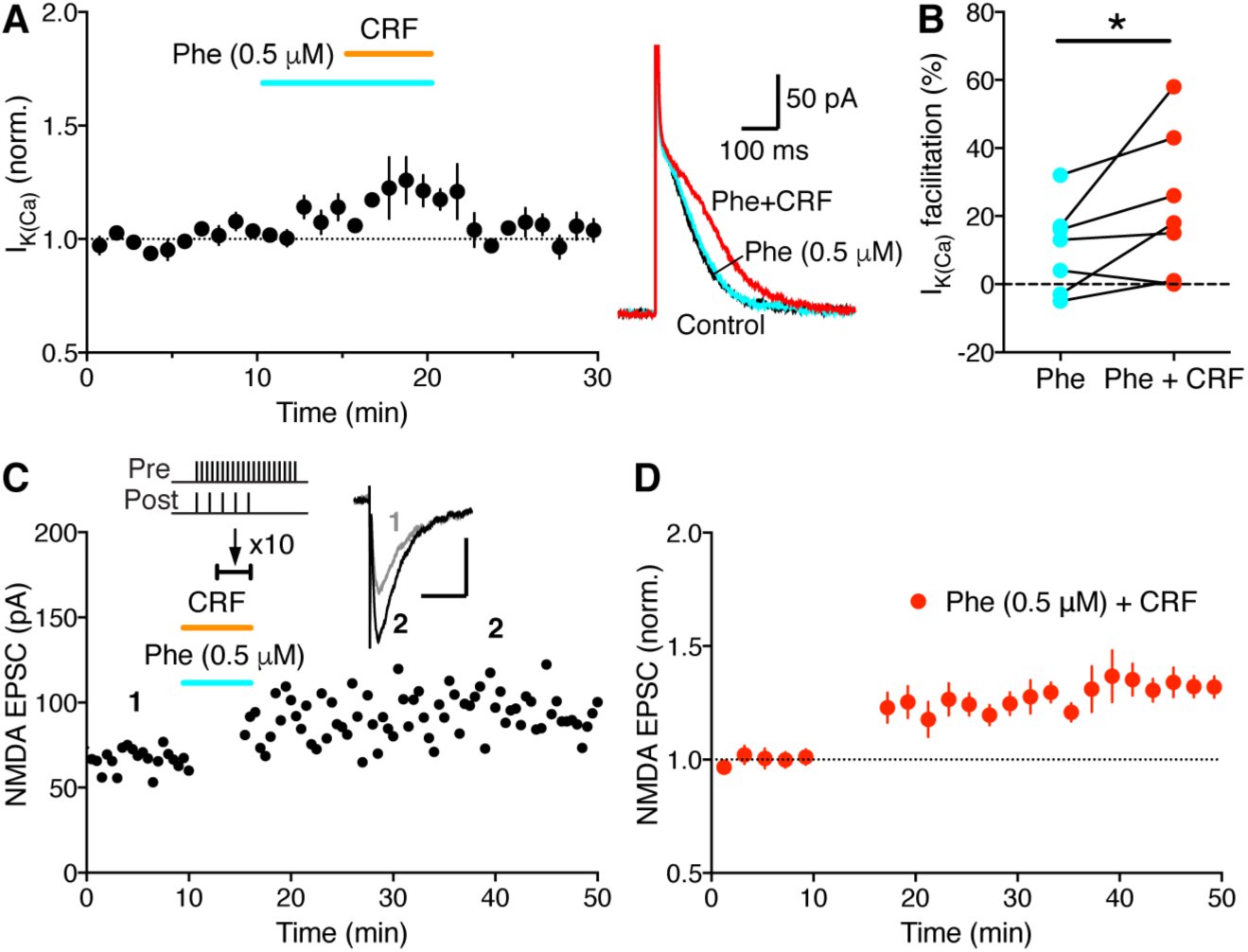
CRF synergizes with phenylephrine to drive LTP without IP_3_-induced Ca^2+^ signal facilitation. (A) Summary time graph (left) and example traces (right) showing that CRF augments facilitation of AP-evoked I_K(Ca)_ produced by a low concentration (0.5 µM) of phenylephrine (7 cells from 5 rats). (B) Graph plotting the magnitude of I_KCa_ facilitation caused by phenylephrine (0.5 µM) alone and by CRF + phenylephrine in individual cells (t_6_ = 2.22, p < 0.05, two-tailed paired t-test). (C) Representative experiment to induce LTP in the presence of both CRF and phenylephrine (0.5 µM) using an induction protocol consisting of synaptic stimulation-burst pairing with no preceding IP_3_ application. Example EPSC traces at the times indicated are shown in inset (scale bars: 50 ms/50 pA). (D) Summary time graph of LTP experiments in which LTP was induced using a synaptic stimulation-burst pairing protocol in the presence of both CRF and phenylephrine (0.5 µM) (7 cells from 4 rats).

Altogether, these data in VTA slices strongly suggest that CRF and NE promote LTP-NMDA by differentially regulating IP_3_-Ca^2+^ signaling, i.e., via CRFR2-mediated increase in IP_3_R sensitivity vs. α1AR-mediated generation of IP_3_, enabling them to act in a synergistic fashion (Figure 6A,B). LTP magnitude was positively correlated with the size of I_K(Ca)_ facilitation during induction across neurons with different induction conditions (Figure 6C), supporting the notion that IP_3_-dependent Ca^2+^ signal facilitation drives LTP.

**Figure 6.**
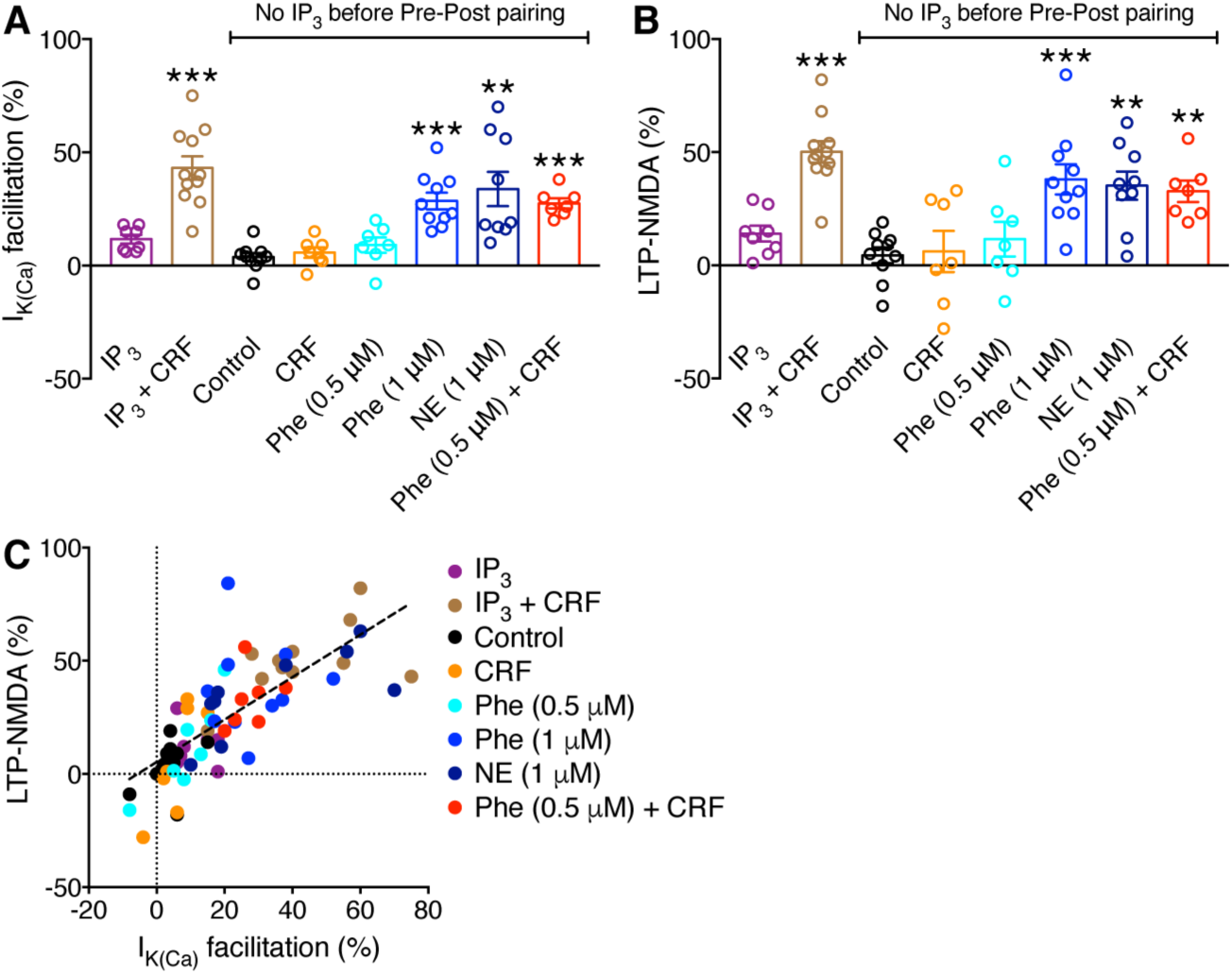
Summary of LTP experiments. (A and B) Summary bar graphs demonstrating the magnitude of I_k(Ca)_ facilitation (A) and LTP (B) for all LTP experiments testing CRF, phenylephrine, and NE (A: F_7,60_ = 13.2, p < 0.0001; B: F_7,60_ = 9.07, p < 0.0001; one-way ANOVA). **p < 0.01, ***p < 0.001 vs. control group with no IP_3_ (Dunnett’s post hoc test). CRF, phenylephrine, and NE had no effect on NMDA transmission itself (Figure 6–figure supplement 1). (C) The magnitude of LTP is plotted versus the magnitude of I_k(Ca)_ facilitation in individual neurons. Dashed line is a linear fit to all data points (n = 69, r^2^ = 0.56).

### CRF and NE synergize in the VTA to drive stress enhancement of cocaine place conditioning

We next sought to explore if CRF and NE actions on NMDA plasticity in the VTA may contribute to social stress-induced enhancement of cocaine CPP illustrated in Figure 1. Low-dose cocaine (5 mg/kg) was used for conditioning in the following experiments to avoid the ceiling effect observed with a higher dose (Fig. 1F). Although delivery of the CRFR2 antagonist K41498 into the VTA prior to social defeat had small effect, stress-enhanced cocaine conditioning was significantly suppressed by the α1AR antagonist prazosin and abolished by co-injection of K41498 and prazosin (Figure 7A–F). Thus acute social defeat stress recruits a cooperative CRF and NE signaling mechanism acting on CRFR2 and α1AR in the VTA to promote cocaine conditioning.

**Figure 7.**
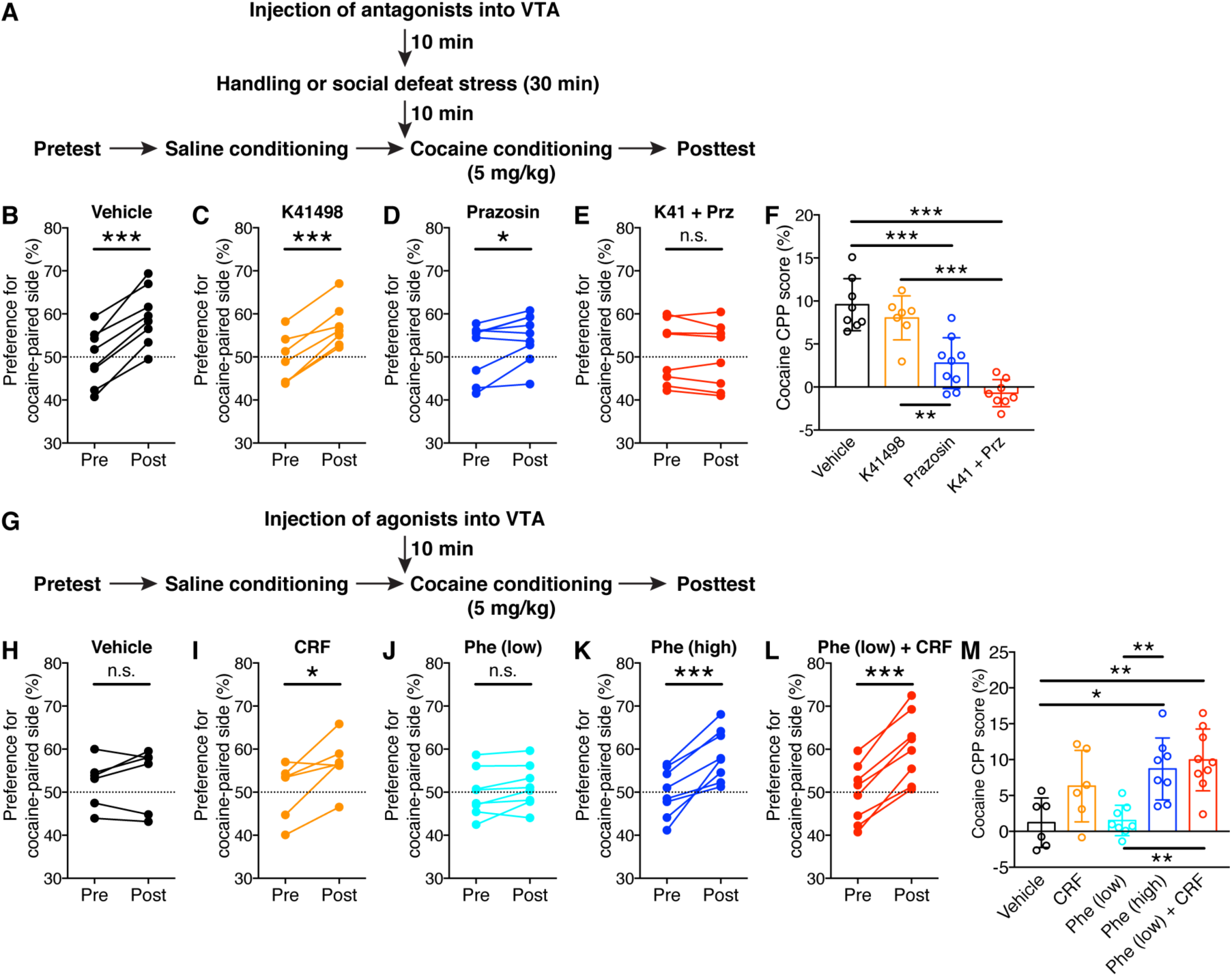
CRF and NE acting on CRFR2 and α1AR in the VTA synergistically promote cocaine place conditioning. (A) Experimental timeline for testing the effects of intra-VTA injection of CRFR2 antagonist K41498 and α1AR antagonist prazosin on defeat stress-induced enhancement of cocaine conditioning. (B–E) Changes in the preference for the cocaine-paired side (conditioned with 5 mg/kg cocaine) in socially defeated rats that received intra-VTA injection of PBS (B), K41498 (C), prazosin (D), or a cocktail of K41498 and prazosin (E) (H: t_7_ = 8.97, p < 0.0001; I: t_7_ = 4.03, p < 0.01; J: t_8_ = 2.82, p < 0.05; K: t_7_ = 1.27, p = 0.24; two-tailed paired t-test). (F) Summary graph demonstrating CRFR2 and α1AR dependence of stress-induced enhancement of cocaine conditioning (F_3,30_ = 14.5, p < 0.0001, one-way ANOVA). **p < 0.01, ***p < 0.001 (Bonferroni post hoc test). (G) Experimental timeline for testing the effects of intra-VTA injection of CRF and phenylephrine on acquisition of cocaine CPP in non-stressed rats.(H–L) Changes in the preference for the cocaine-paired side (conditioned with 5 mg/kg cocaine) in rats that received intra-VTA injection of PBS (H), CRF (I), high-dose phenylephrine (18 pmol/0.3 µL; J), low-dose phenylephrine (6 pmol/0.3 µL; K), or a cocktail of CRF and low-dose phenylephrine (L) (B: t_7_ = 0.40, p = 0.70; C: t_7_ = 2.28, p = 0.057; D: t_7_ = 5.69, p < 0.001; E: t_7_ = 2.02, p = 0.083; F: t_7_ = 8.89, p < 0.0001; two-tailed paired t-test). (M) Summary graph demonstrating the effects of CRF and phenylephrine on cocaine place conditioning in the absence of stress (F_4,36_ = 5.17, p < 0.01, one-way ANOVA). *p < 0.05, **p < 0.01 (Bonferroni post hoc test).

Are CRF and NE actions in the VTA sufficient to enhance cocaine conditioning in the absence of stress (Figure 7G)? While control rats injected with vehicle (PBS) into the VTA developed inconsistent CPP, intra-VTA microinjection of CRF (1.5 pmol/0.3 µL/side) prior to cocaine conditioning enabled moderate CPP (Figure 7H,I,M). We further found that administration of the α1AR agonist phenylephrine (18 pmol/0.3 µL/side) lead to robust cocaine conditioning, although a lower dose (6 pmol/0.3 µL/side) had minimal effect (Figure 7J,K,M). Notably, combined application of CRF with low-dose phenylephrine enabled large CPP comparable to that observed with high-dose phenylephrine (Figure 7L,M). These data further support the idea that CRF and NE synergize in the VTA to enhance cocaine conditioning.

### Acute social defeat stress promotes Pavlovian cue-food conditioning and alleviates temporal constraints on learning

LTP-NMDA in DA neurons is induced in a burst-timing-dependent manner, where the onset of glutamatergic input stimulation needs to precede postsynaptic burst, reflecting the kinetics of mGluR-induced rise in IP_3_ to reach effective levels at the time of burst (Harnett et al., 2009). This timing dependence of LTP might partially account for the need of a delay between cue onset and reward delivery for the acquisition of cue-evoked DA neuron bursting (Cohen et al., 2012; Kobayashi and Schultz, 2008) and appetitive learning (Pavlov, 1927; Schwartz et al., 2002). If so, acute stress might alter the cue-reward timing rule via NE action generating IP_3_, boosted by CRF effect on IP_3_R sensitivity. To test this idea, we used a Pavlovian conditioned approach paradigm and varied the temporal relationship between the onset of cue (10 sec light at food magazine) and delivery of reward (food pellet) during conditioning (Figure 8A and Figure 8–figure supplement 1). While both handled controls and defeated rats developed a comparable conditioned response to the food-paired cue (magazine entry, i.e., approach to the cue light) after one conditioning session with 5 sec cue-reward delay (Figure 8B,C and Figure 8–figure supplement 2), only defeated rats developed a cue response when the cue onset and reward delivery were simultaneous (Figure 8D,E and Figure 8–figure supplement 2). Neither group developed a conditioned response when the reward was delivered 5 sec prior to cue onset (Figure 8F,G and Figure 8–figure supplement 2), suggesting that cue-encoding neural activity needs to be active at the time of reward. Thus acute defeat stress appeared to shift the cue-reward timing dependence, minimizing the requirement of delay to drive effective conditioning (Figure 8H).

**Figure 8.**
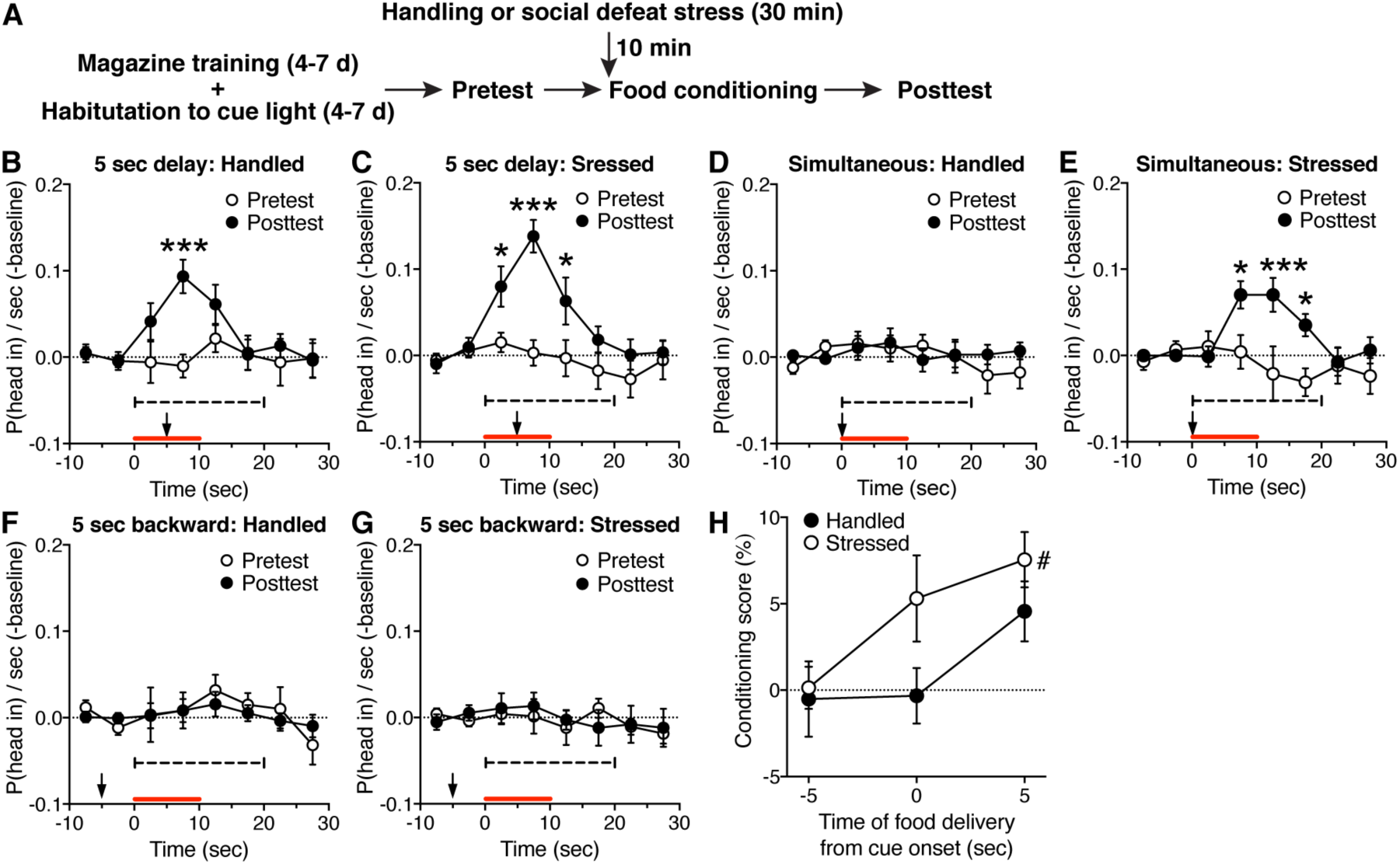
Acute social defeat stress enables learning of a food-paired cue with no delay between cue onset and food delivery. (A) Experimental timeline for testing the effect of acute social defeat stress on food conditioned approach. (B–G) Summary time graphs showing cue responses (binned in 5 sec) before and after conditioning in handled and defeated rats (B: 9 rats, C: 11 rats, D: 12 rats, E: 11 rats, F: 9 rats, G: 10 rats). Cue light was presented at the red bar, while the arrow indicates the time of food delivery during conditioning (B: time: F_7,56_ = 3.30, p < 0.01; time × test: F_7,56_ = 2.31, p < 0.05; C: time: F_7,70_ = 7.22, p < 0.0001; test: F_1,10_ = 19.2, p < 0.01; time × test: F_7,70_ = 4.81, p < 0.001; E: time: F_7,70_ = 3.34, p < 0.01; time × test: F_7,56_ = 3.30, p < 0.01; repeated measures two-way ANOVA). (H) Summary graph illustrating the cue-reward timing dependence of conditioning in control and stressed rats (time of food: F_2,56_ = 5.33, p < 0.01; stress: F_1,56_ = 4.09, p < 0.05; two-way ANOVA). Conditioning score (expressed in %) was calculated from the 20 sec period at the dashed line in (B)–(G). *p < 0.05, ***p < 0.001 vs. pretest in (B), (C), and (E); ^#^p < 0.05 vs. -5 sec group in (H) (Bonferroni post hoc test).

## DISCUSSION

Multiple stress mediators, including glucocorticoids acting via a rapid non-genomic pathway, interact, sometimes in an antagonistic fashion, to acutely regulate synaptic plasticity and learning and memory processes (Joels et al., 2011; Maras and Baram, 2012; McEwen, 2007). For example, corticosterone can promote or suppress the facilitatory effect of NE, acting via β adrenergic receptors (βARs), on synaptic plasticity depending on the timing of application in the hippocampus and amygdala (Akirav and Richter-Levin, 2002; Pu et al., 2007, 2009), while a recent study reported a synergistic action of corticosterone and CRF to impair hippocampal glutamatergic synapses and spatial memory (Chen et al., 2016). The present study demonstrates that CRF and NE synergistically augment IP-Ca2+ signaling, via CRFR2-dependent increase in IP_3_ sensitivity and α1AR-dependent IP3 synthesis, respectively, driving enhanced plasticity of NMDA transmission in VTA DA neurons. Our data further implicate a synergistic action of CRFR2 and α1AR signaling in acute social defeat stress-induced enhancement of cocaine place conditioning. Thus this study identifies a potential molecular target on which the two acute stress mediators act in concert to regulate a form of appetitive learning.

While previous studies reporting CRF/NE-induced enhancement of AMPA plasticity have mostly focused on regulation of neuronal excitability (Blank et al., 2002; Liu et al., 2017) or postsynaptic AMPA receptors (Hu et al., 2007; Seol et al., 2007), our study implicates CRF/NE effects on a Ca^2+^-dependent induction process per se as the metaplasticity mechanism for NMDA plasticity. Interestingly, NE acting on βARs has been shown to enhance spike-timing-dependent plasticity in the hippocampus by relieving the constraints on the timing of pre- and postsynaptic spikes (Lin et al., 2003; Seol et al., 2007) or on the number of postsynaptic spikes (Liu et al., 2017). The current study suggests that NE acting on α1ARs to generate IP_3_, together with CRF facilitating this α1AR effect, may remove the requirement of presynaptic stimulation preceding postsynaptic bursting. Thus stress mediators appear to lower the “gate” for synaptic plasticity at multiple levels in different brain areas.

Although our study has identified a critical role of CRFR2 in the VTA in promoting NMDA plasticity and cocaine conditioning, it is known that DA neurons also express CRFR1, which can control DA neuron physiology and reward/drug-driven behaviors (Henckens et al., 2016). For example, while no significant effect of CRF (100– 300 nM) on NMDA transmission was observed in the current study, previous studies have reported CRF effects on NMDA and AMPA transmission in VTA DA neurons, involving multiple mechanisms via both CRFR1 and CRFR2 depending on the CRF concentration used (Hahn et al., 2009; Ungless et al., 2003; Williams et al., 2014). It remains to be determined how multiple CRF effects on glutamatergic transmission reflect heterogeneity of DA neurons in the VTA, especially given differential effects of appetitive vs. aversive/stressful stimuli on these neurons (Holly and Miczek, 2016; Lammel et al., 2011; Morales and Margolis, 2017; Polter and Kauer, 2014). Regardless, these CRFR1/CRFR2-dependent effects on glutamatergic excitation, together with CRF/NE effects on DA neuron firing (Paladini et al., 2001; Wanat et al., 2008), may contribute to the acute stress-induced enhancement of the expression of drug-seeking behavior observed in vivo (Holly et al., 2016; Mantsch et al., 2016; Wang et al., 2007). It should be noted that CRF and NE actions in other limbic structures also contribute to different aspects of reward-driven behavior (Henckens et al., 2016; Otis et al., 2015; Smith and Aston-Jones, 2008). Despite the engagement of multiple brain circuits in response to acute stress-induced CRF/NE actions, our data implicate VTA DA neuron plasticity as the critical substrate for enhancement of appetitive cue learning.

The VTA receives inputs from several CRF-rich regions including the bed nucleus of the stria terminalis, central amygdala, and paraventricular hypothalamus, and paraventricular hypothalamus (Rodaros et al., 2007), while major sources of NE to the VTA are the locus coeruleus and A1, A2, and A5 noradrenergic cell groups that exhibit distinct topography of innervation to VTA subareas (Mejias-Aponte et al., 2009). Indeed, many of these brain areas are activated by social defeat stress (Martinez et al., 1998). Different types of stress may differentially recruit CRF and NE sources to the VTA, thus creating different levels of CRF and NE to regulate their synergistic interaction.

It is well known that brief stressful experience could lead to persistent changes in brain function depending on the intensity or controllability of the stressor (Musazzi et al., 2017). Indeed, a number of studies have shown persistent changes in VTA synapses lasting >1 day following single or repeated stress exposure, which are frequently linked to intensification and/or reinstatement of drug-seeking behavior (Polter and Kauer, 2014). Our previous study has shown that repeated (5 day), but not single, social defeat stress promotes the NMDA plasticity mechanism 1-10 days later, which is associated with enhanced cocaine CPP (Stelly et al., 2016). Enhancement of plasticity and CPP both require glucocorticoid action during stress exposure, likely through glucocorticoid receptors mediating long-lasting changes in gene expression. Although blockade of CRF and NE signaling in the VTA completely suppressed acute stress effect on CPP in the current study, it may be possible that rapid non-genomic glucocorticoid effects may play a permissive role, as has been demonstrated for the effects of CRF and/or NE on synaptic function and memory processes in the hippocampus and amygdala (Chen et al., 2016; Roozendaal et al., 2008).

Interestingly, acute stress (inescapable electric shock or swim stress) has been shown to enhance Pavlovian eyeblink conditioning (Shors, 2001; Shors et al., 1992), which may be driven by a form of synaptic plasticity in the cerebellum that is dependent on an IP_3_-Ca^2+^ signaling mechanism similar to NMDA plasticity in DA neurons (Wang et al., 2000). This facilitatory effect on eyeblink conditioning can be observed 30 min to 24 hr after stress exposure, while the effect on cocaine CPP was observed 30 min, but not 1.5 hr (current study) or 24 hr (Stelly et al., 2016), following a single episode of defeat stress. The role of different stress mediators underlying the persistence of single stress exposure on eyeblink conditioning has not been explored, although the effects of CRF and NE on cerebellar synaptic plasticity have been reported (Carey and Regehr, 2009; Schmolesky et al., 2007). It should also be noted that a single exposure to inescapable footshock or restraint stress has been reported to promote CPP acquisition for days (Pacchioni et al., 2002; Will et al., 1998).

In the present study, the facilitatory effect of acute social defeat stress on Pavlovian cue learning was observed not only with cocaine (i.e., drug reward) but also when food reward was used as an unconditioned stimulus (US) for conditioning. Indeed, a recent human study has reported that brief exposure to cold stress 2 min prior to Pavlovian conditioning sessions using monetary rewards promoted cue-evoked activity in the ventral striatum (Lewis et al., 2014). It is well established that the cue needs to be presented prior to the US for different types of Pavlovian conditioning (Pavlov, 1927; Schwartz et al., 2002). Intriguingly, stressed rats acquired a cue response (i.e., approach to the cue light) even when the cue and US (food pellet) were presented simultaneously, in apparent violation of a canonical principle of Pavlovian conditioning. Although speculative, concerted CRF and NE actions on IP_3_ signaling in VTA DA neurons might mitigate the requirement of the delay from cue onset to reward delivery, during which cue-evoked glutamatergic input activating mGluRs is hypothesized to cause IP_3_ rise at the time of reward-evoked bursting to effectively drive NMDA potentiation (Harnett et al., 2009). By enabling simultaneous cue-reward conditioning, daily stressful experience may lead to spurious learning of reward-associated cues with no predictive value or redundant cues presented at the same time with already learned cues (e.g., money) (Holland, 1984), thereby driving increased addiction liability to drug and non-drug rewards.

## METHODS AND MATERIALS

### Animals

Sprague-Dawley rats (Harlan Laboratories, Houston, Texas) were housed in pairs on a 12-hr light/dark cycle with food and water available ad libitum. All procedures were approved by the University of Texas Institutional Animal Care and Use Committee.

### Brain slice electrophysiology

Midbrain slices were prepared and recordings were made in the lateral VTA located 50– 150 mm from the medial border of the medial terminal nucleus of the accessory optic tract, as in our previous studies (Stelly et al., 2016; Whitaker et al., 2013). Tyrosine hydroxylase-positive neurons in this area (i.e., lateral part of the parabrachial pigmented nucleus) largely project to the ventrolateral striatum (Ikemoto, 2007) and show little VGluT2 coexpression (Trudeau et al., 2014). Internal solution contained (in mM): 115 K-methylsulfate, 20 KCl, 1.5 MgCl_2_, 10 HEPES, 0.025 EGTA, 2 Mg-ATP, 0.2 Na_2_-GTP,and 10 Na_2_-phosphocreatine (pH ∼7.25, ∼285 mOsm/kg). Putative dopamine neurons in the lateral VTA were identified by spontaneous firing of broad APs (>1.2 ms) at 1–5 Hz in cell-attached configuration and large I_h_ currents (>200 pA; evoked by a 1.5 s hyperpolarizing step of 50 mV) in whole-cell configuration (Ford et al., 2006; Lammel et al., 2008; Margolis et al., 2008). Cells were voltage-clamped at –62 mV (corrected for –7 mV liquid junction potential). A 2 ms depolarizing pulse of 55 mV was used to elicit an unclamped AP. For bursts, 5 APs were evoked at 20 Hz. The time integral of the outward tail current, termed I_K(Ca)_ (calculated after removing the 20 ms window following each depolarizing pulse; expressed in pC), was used as a readout of AP-evoked Ca^2+^ transients, as it is eliminated by TTX and also by apamin, a blocker of Ca^2+^-activated SK channels (Cui et al., 2007).

### UV Photolysis

Cells were loaded with caged IP_3_ (1–10 µM) through the recording pipette. UV light (100 ms) was applied using the excitation light from the xenon arc lamp of the Olympus Disk Spinning Unit imaging system. The light was focused through a 60× objective onto a ∼350 µm area surrounding the recorded neuron. Photolysis of caged compounds is proportional to the UV light intensity, which was adjusted with neutral density filters and measured at the focal plane of the objective (in mW). The applied IP_3_ concentration is expressed in µM·mW.

### LTP experiments

Synaptic stimuli were delivered with a bipolar tungsten electrode placed ∼200 µm rostral to the recorded neuron. To isolate NMDA EPSCs, recordings were performed in DNQX (10 µM), picrotoxin (100 µM), CGP54626 (50 nM), and sulpiride (100 nM) to block AMPA/kainate, GABA_A_, GABA_B_, and D_2_ dopamine receptors, and in glycine (20 µM) and low Mg^2+^ (0.1 mM) to enhance NMDA receptor activation. NMDA EPSCs were monitored every 20 s. The LTP induction protocol consisted of photolytic application of IP_3_ (10 µM·mW) for 100 ms prior to the simultaneous delivery of afferent stimulation (20 stimuli at 50 Hz) and postsynaptic burst (5 APs at 20 Hz), repeated 10 times every 20 s. LTP magnitude was determined by comparing the average EPSC amplitude 30-40 min post-induction with the average EPSC amplitude pre-induction (each from a 10 min window).

### Resident-Intruder Social Defeat Paradigm

Twelve week-old male resident rats were vasectomized and pair-housed with 6 week-old females. Residents (used for ∼8–10 months) were screened for aggression (biting or pinning within 1 min) by introducing a male intruder to the home cage. Intruders and controls were young males (4–5 weeks old at the beginning) housed in pairs. For defeat sessions, intruders were introduced to residents’ home cages after removing females. Following ∼5 min of direct contact, a perforated Plexiglass barrier was inserted for ∼25 min to allow sensory contact, as in our previous study (Stelly et al., 2016). The barrier was removed for a brief period (<1 min) in certain cases to encourage residents’ threatening behavior. Handled controls were placed in novel cages for 30 min. Intruders and controls were housed separately.

### Cocaine Place Conditioning

CPP boxes (Med Associates) consisting of two distinct compartments separated by a small middle chamber were used for conditioning. One compartment had a mesh floor with white walls, while the other had a grid floor with black walls. A discrete cue (painted ceramic weight) was placed in the rear corner of each compartment (black one in the white wall side, white one in the black wall side) for further differentiation. Rats were first subjected to a pretest, in which they explored the entire CPP box for 15 min. The percentage of time spent in each compartment was determined after excluding the time spent in the middle chamber. Rats with initial side preference >60% were excluded. The following day, rats were given a saline injection in the morning and confined to one compartment, then in the afternoon given cocaine (5 or 10 mg/kg, i.p.) and confined to the other compartment (10 min each). Compartment assignment was counterbalanced such that animals had, on average, ∼50% initial preference for the cocaine-paired side. A 15 min posttest was performed 1 day after conditioning. The CPP score was determined by subtracting the preference for the cocaine-paired side during pretest from that during posttest. The experimenter performing CPP experiments was blind to animal treatments.

### Intra-VTA microinjections

Rats (7–10 weeks old) were anesthetized with a mixture of ketamine and xylazine (90 mg/kg and 10 mg/kg, i.p.) and implanted with bilateral chronic guide cannulas (22 gauge; Plastics One), with dummy cannulas (32 gauge) inside, aimed at 1 mm above the VTA (anteroposterior, –5.3; mediolateral, +2.2; dorsoventral, –7.5; 10° angle). The guide cannulas were fixed to the skull with stainless steal screws and dental cement. After the surgery, rats remained singly housed for 7 days before being subjected to conditioning experiments.

Intra-VTA microinjections were made via injection cannulas (28 gauge; Plastics One) that extended 1 mm beyond the tip of the guide cannulas. Injection cannulas were connected to 1 µL Hamilton syringes mounted on a microdrive pump (Harvard apparatus). Rats received bilateral infusions (0.3 µL/side, 0.15 µL/min) of different pharmacological agents in certain conditioning experiments. The injection cannulas were left in place for 60 s after infusion.

At the end of conditioning experiments, rats were anesthetized with a mixture of ketamine and xylazine (90 mg/kg and 10 mg/kg, i.p.) and transcardially perfused with 4% paraformaldehyde. Brains were then carefully removed and stored in 4% paraformaldehyde. Coronal sections (100 µm) were cut using a vibratome and stained with cresyl violet for histological verification of injections sites (Figure 7–figure supplement 1). Data from rats with injection sites outside the VTA were excluded from the analysis.

### Pavlovian Conditioned Approach

Conditioning was performed in modular test chambers (Med Associates) equipped with a food pellet receptacle at the center of one wall. Illuminating light at the rear of receptacle was used as a cue during conditioning (house light was turned off). Head entry was detected with infrared photobeam positioned across the receptacle. All sessions were performed on a 60 sec variable inter-trial interval schedule (range 40–80 sec). Each session was preceded by a 5 min acclimation period during which rats stayed in the chamber with no food pellet delivery or cue light illumination. Rats first underwent 4–7 days of magazine training sessions in which rats received 30 banana-flavored food pellets (45 mg; Bio-Serv) with no light illumination at the receptacle and learn to rapidly (within ∼1 sec) respond to the food drop sound (Figure 8–figure supplement 3). To minimize unconditioned response to the receptacle light, rats received 4–7 days of habituation sessions where rats were exposed to 10 sec illumination of receptacle light with no food delivery (15–30 trials per day; alternated with several days of magazine training sessions). The final habituation session (15 trials) was used as a pretest to assess the response to the cue light before conditioning. On the day following this pretest, rats underwent 30 trial conditioning sessions, in which the food pellet was delivered either at the onset of the 10 sec light cue, 5 sec after the cue onset (i.e., at the middle of 10 sec light cue), or 5 sec before the cue onset. Posttest (15 trials, cue light with no food) was performed 1 day after conditioning. Responses were measured with the proportion of trials in which head entry was detected in each second (labeled P(head in)/sec). The mean value during the 10 sec baseline period before cue onset was subtracted in each rat to assess the cue light response. Rats displaying significant non-habituated cue response during the pretest (mean P(head in)/sec >0.1 above baseline level during the 20 sec period from cue onset, averaged over 15 trials) were excluded from analysis. The conditioning score was determined by subtracting the mean P(head in)/sec above baseline level in the pretest from that in the posttest (expressed in %).

### Drugs

DNQX, picrotoxin, CGP55845, sulpiride, CRF and K41498 were obtained from Tocris Biosciences. Caged IP3 was a generous gift from Dr. Kamran Khodakhah (Albert Einstein College of Medicine). All other chemicals were from Sigma-RBI.

### Data Analysis

Data are expressed as mean ± SEM. Statistical significance was determined by Student’s t test or ANOVA followed by Bonferroni or Dunnett’s post hoc test. The difference was considered significant at p < 0.05.

## ACKNOWLEDGEMENTS

We thank Dr. Kamran Khodakhah for the generous gift of caged IP_3_ made in his lab at Albert Einstein College of Medicine. We also thank Dr. Claire Stelly for comments on this manuscript.

## FIGURE SUPPLEMENTS

**Figure 2–figure supplement 1.**
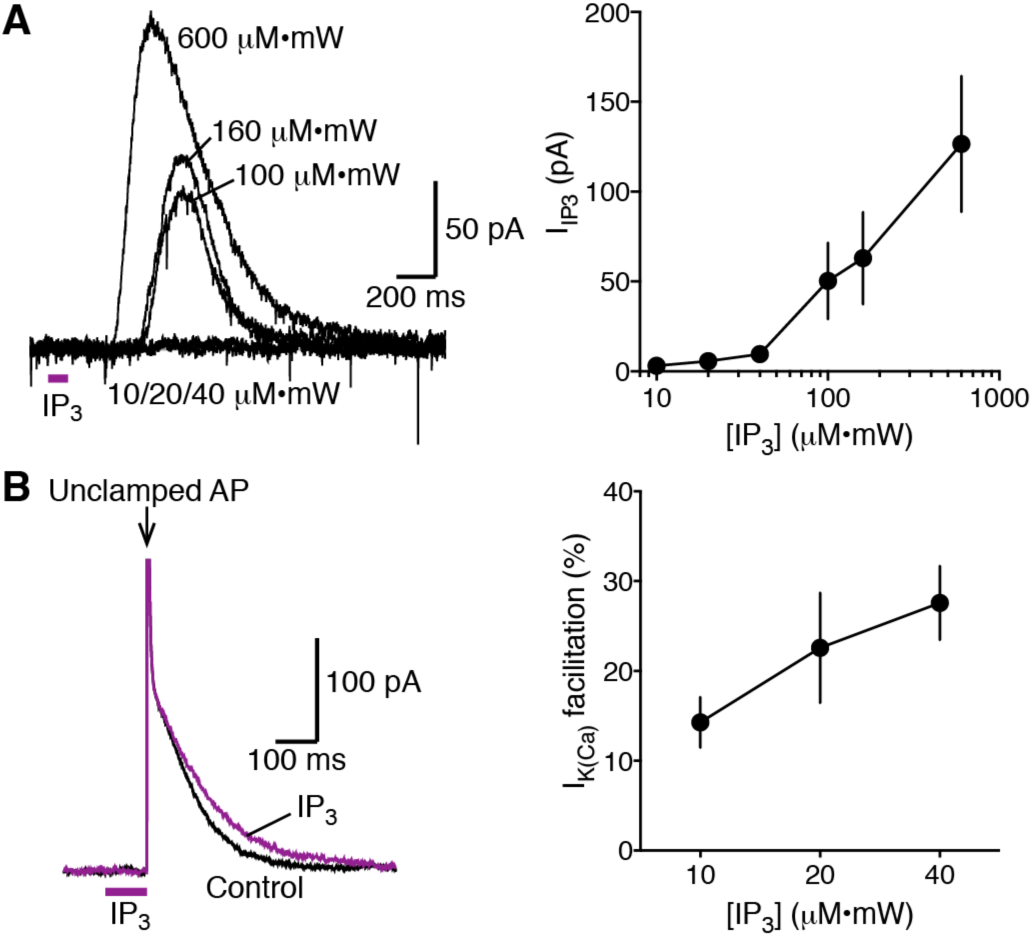
(A) Example traces and summary graph depicting the concentration dependence of IP_3_-evoked outward currents. Data were obtained from 7 cells, where six different IP_3_ concentrations (10, 20, 40, 100, 160, and 600 µM·mW; photolytically applied for 100 ms) were tested in each cell. (B) Example traces (using 40 µM·mW IP_3_) and summary graph illustrating facilitation of AP-evoked I_K(Ca)_ caused by near-threshold levels of IP_3_ (10, 20, and 40 µM·mW; n = 14, 7, and 7, respectively). Note the relatively long latency (∼200–400 ms) following IP_3_ application to evoke response at suprathreshold range (100, 160, and 600 µM·mW), which reflects the time required to engage the regenerative IP_3_R-mediated Ca^2+^-induced Ca^2+^ release process. In contrast, IP_3_ effect on AP-evoked I_K(Ca)_ occurs with no latency, as rapid Ca^2+^ influx triggered by APs initiates the Ca^2+^-induced Ca^2+^ release process, which can be augmented by near-threshold levels of IP_3_.

**Figure 4–figure supplement 1.**
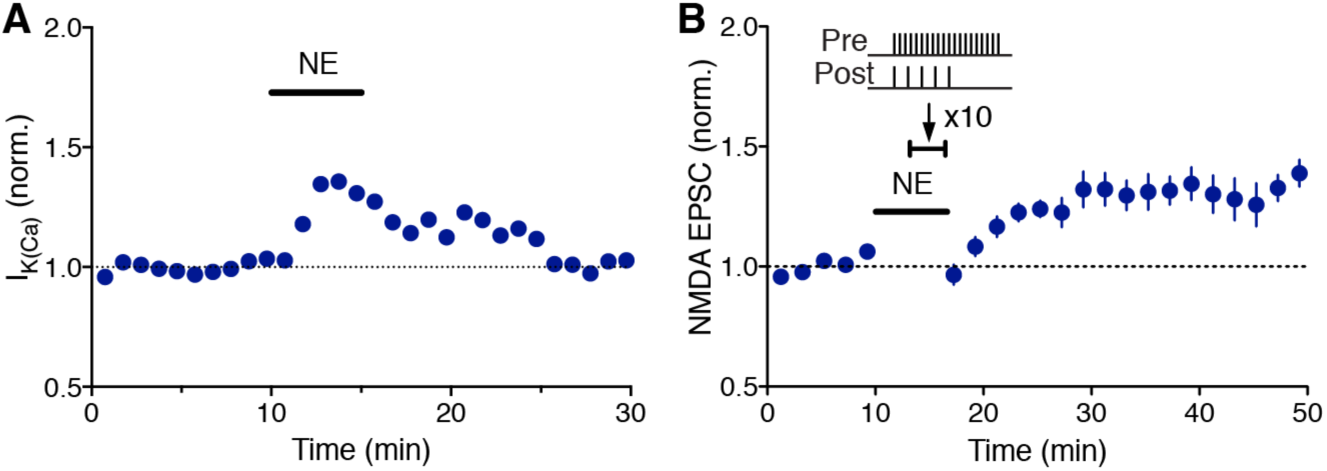
(A) Summary time graph showing the facilitatory effect of NE (1 µM) on AP-evoked I_K(Ca)_ (n = 5). (B) Summary time graph of LTP-NMDA experiments in which LTP was induced using a synaptic stimulation-burst pairing protocol in the presence of NE (1 µM; n = 9).

**Figure 5–figure supplement 1.**
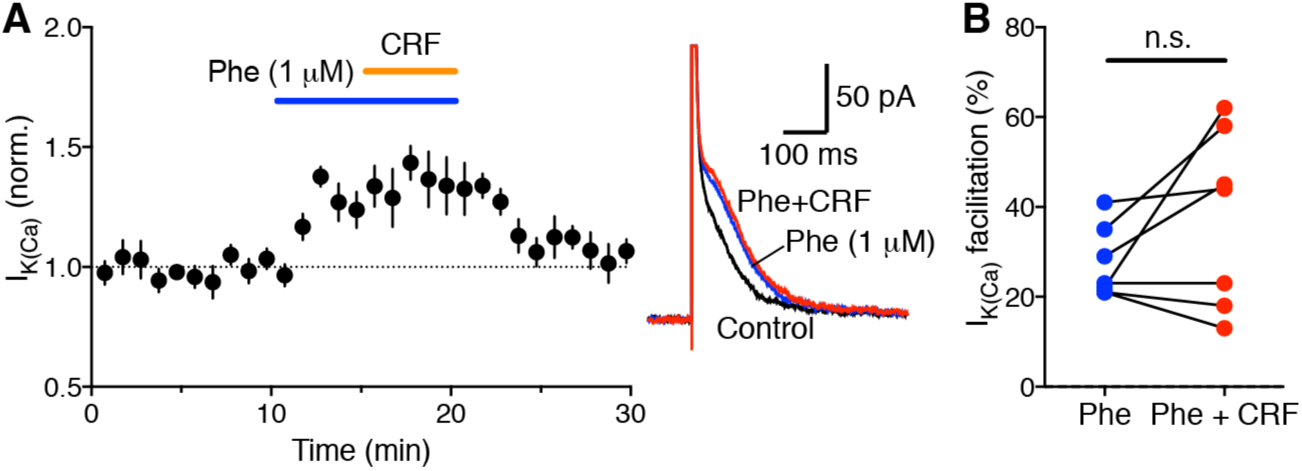
(A) Summary time graph (left) and example traces (right) showing that CRF does not have significant effect on AP-evoked IK(Ca) facilitated by a high concentration (1 µM) of phenylephrine (n = 9). (B) Graph plotting the magnitude of IK(Ca) facilitation caused by phenylephrine (1 µM) alone and by CRF + phenylephrine in individual cells (t_6_ = 1.57, p = 0.17, two-tailed paired t-test).

**Figure 6–figure supplement 1.**
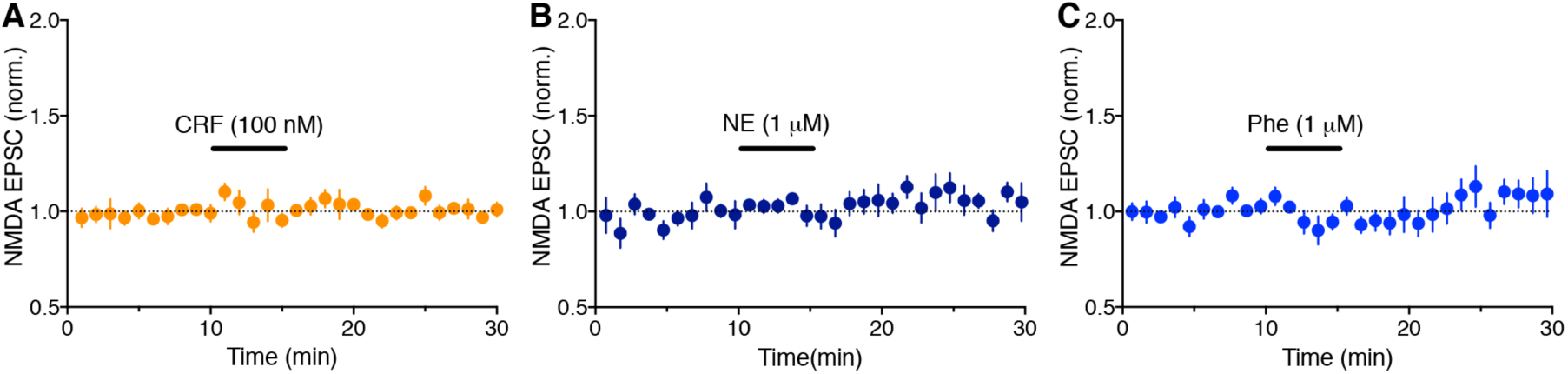
Summary time graphs showing that CRF (A: n = 5), NE (B: n = 7), and phenylephrine (C: n = 5) have no measurable effect on NMDA EPSCs.

**Figure 7–figure supplement 1.**
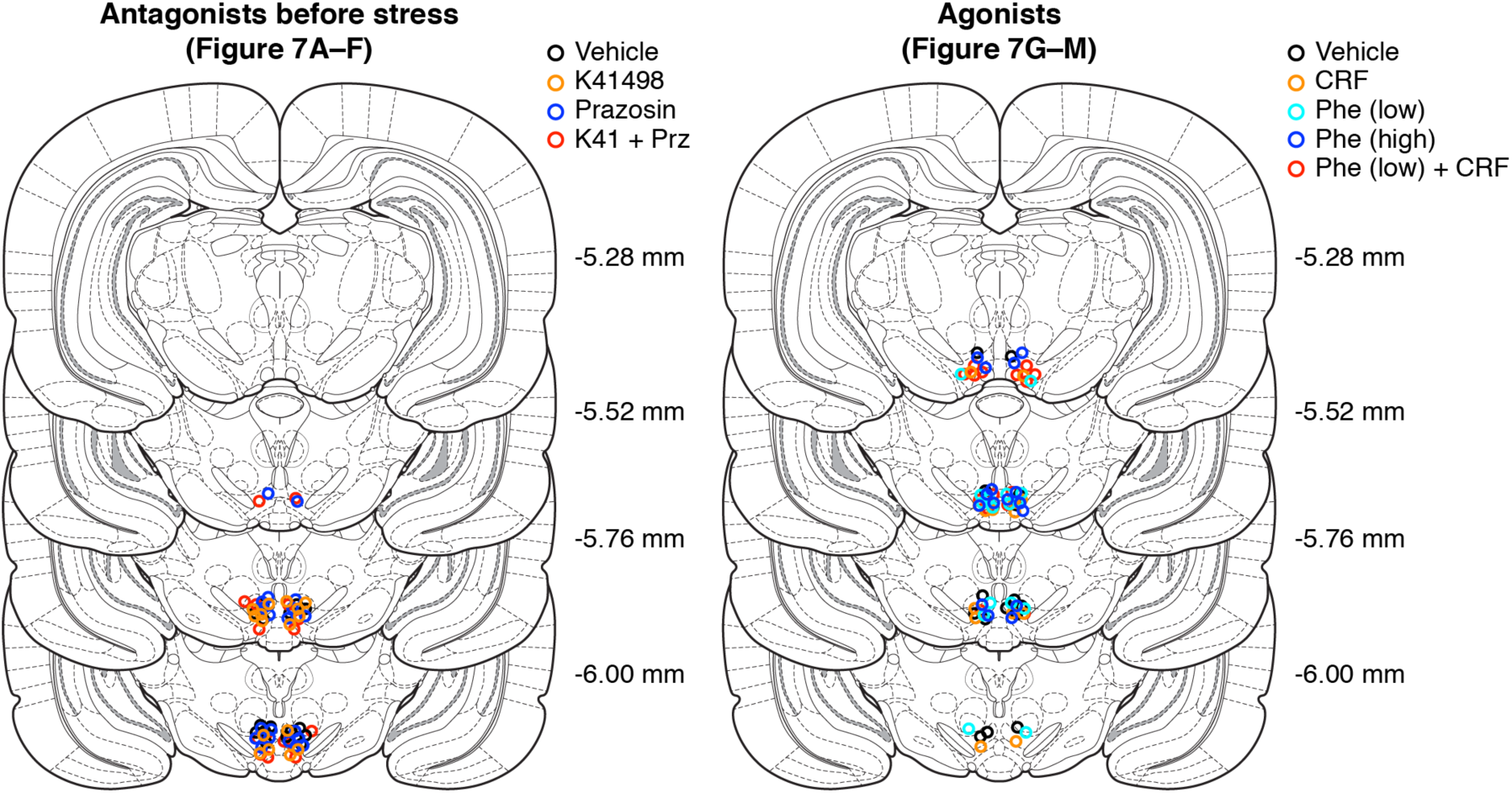
Approximate locations (mm from bregma) of cannula tips for intra-VTA microinjection experiments in Figure 7.

**Figure 8–figure supplement 1.**
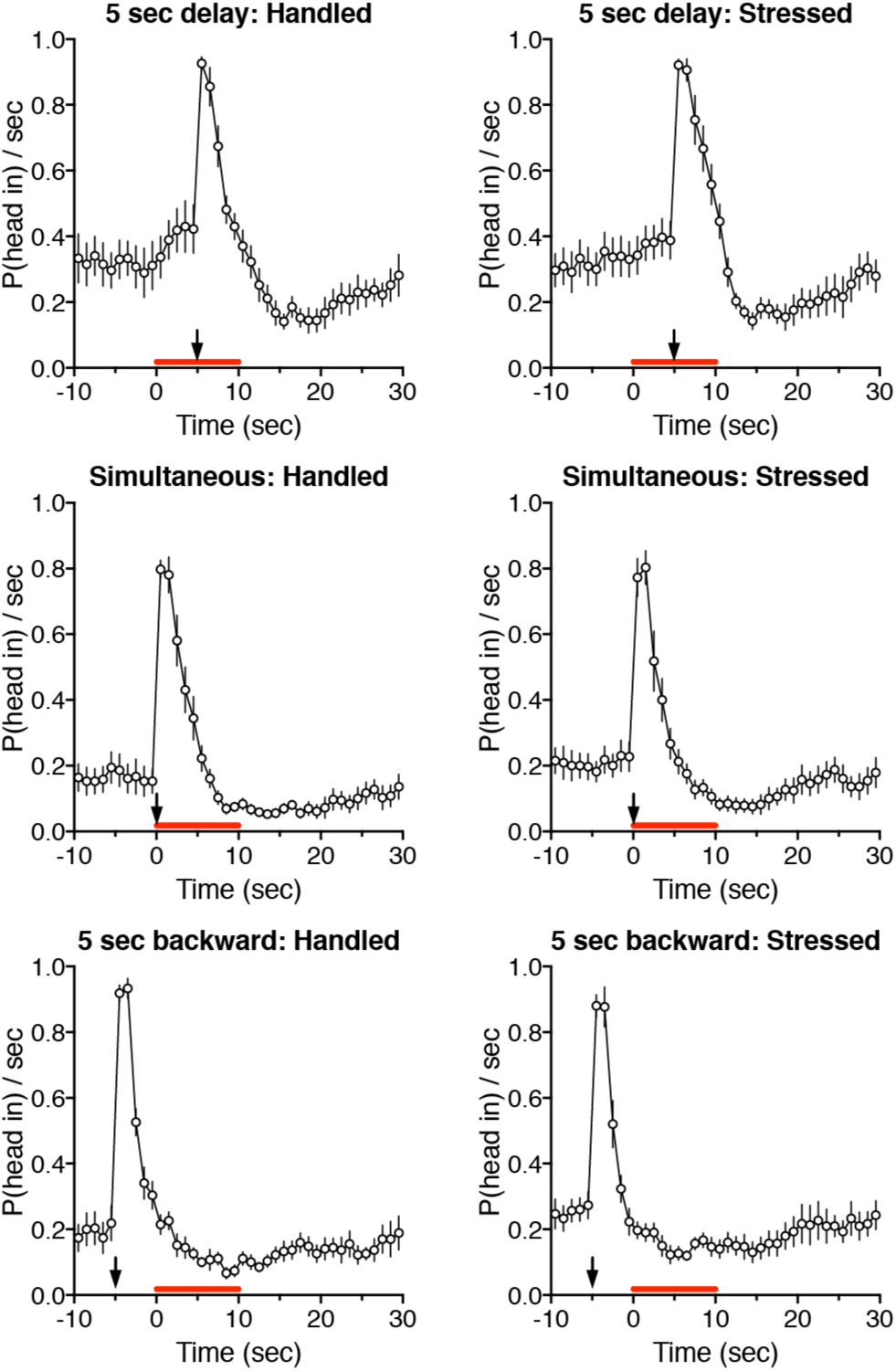
Time graphs illustrating head entry responses during conditioning sessions. The 10 sec cue light was presented at the red bar, while food was delivered at arrow.

**Figure 8–figure supplement 2.**
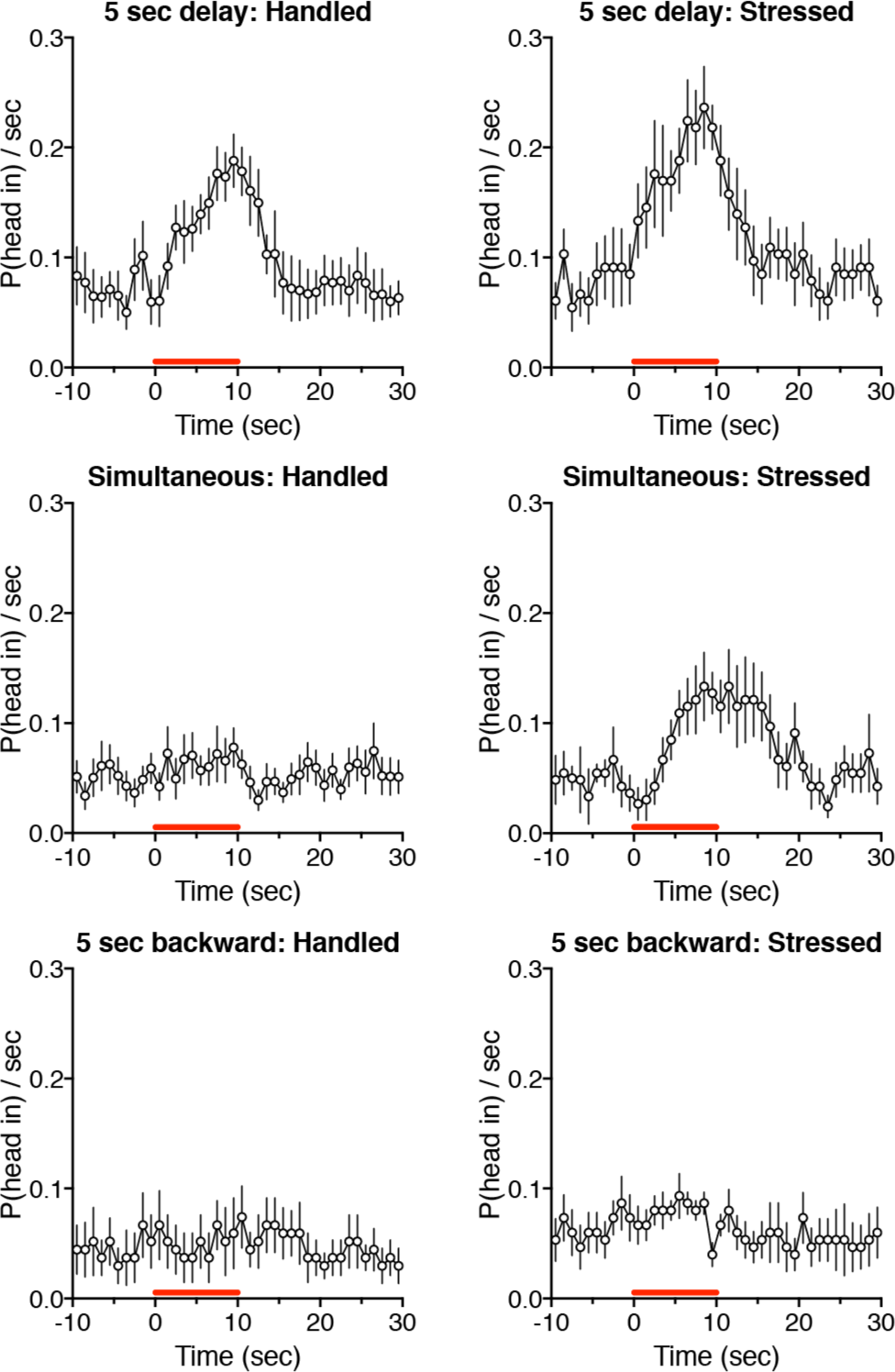
Time graphs illustrating head entry responses during posttests. The 10 sec cue light was presented at the red bar.

**Figure 8–figure supplement 3.**
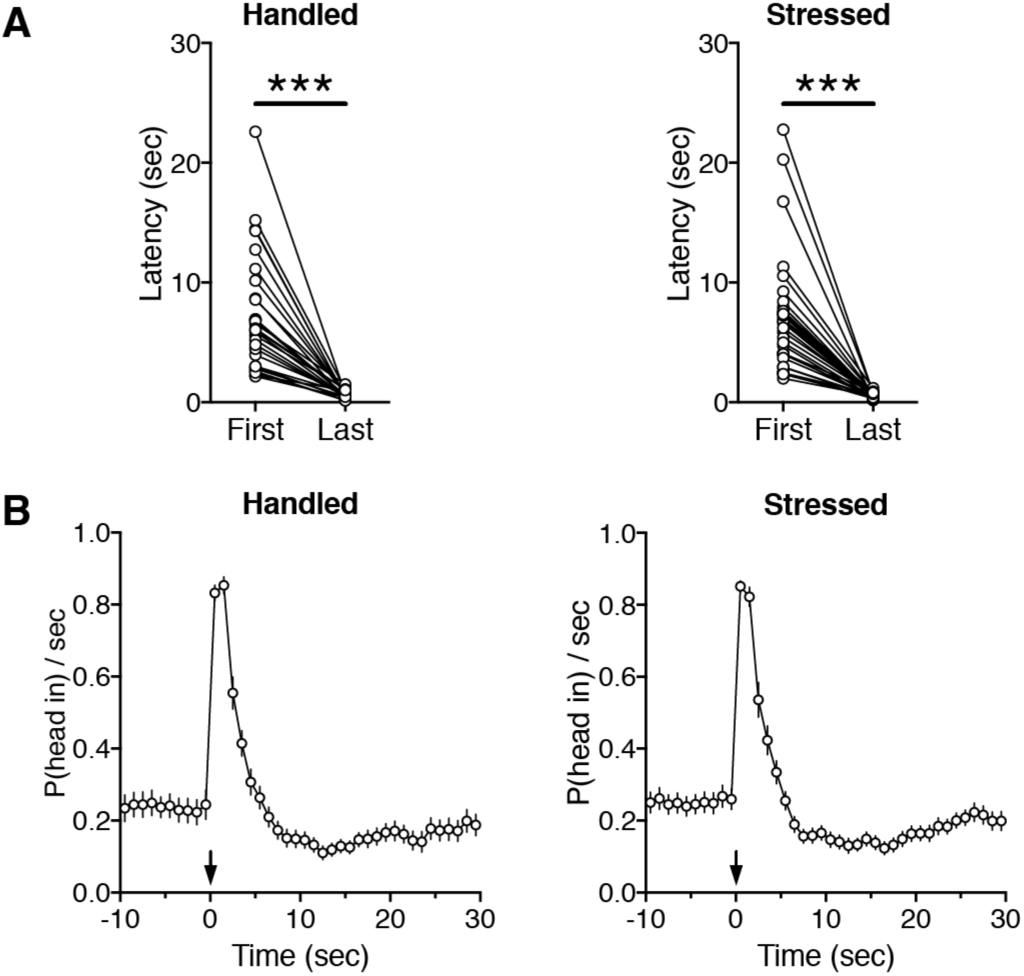
Graphs depicting head entry responses during magazine training sessions before undergoing handling/social defeat and conditioning sessions. (A) Mean latency to the first head entry after food delivery is plotted during the first and last magazine training sessions in individual rats. Data are from all rats shown in Figure 8 (handled: t_29_ = 7.52, p < 0.0001; stressed: t_31_ = 7.53, p < 0.0001; paired t-test). (B) Time graphs plotting the probability of head entry into the food magazine during the last magazine training session. Food was delivered at arrow.

